# Differential protein expression marks the transition from infection with *Opisthorchis viverrini* to cholangiocarcinoma

**DOI:** 10.1101/086645

**Authors:** Jarinya Khoontawad, Chawalit Pairojkul, Rucksak Rucksaken, Porntip Pinlaor, Chaisiri Wongkham, Puangrat Yongvanit, Ake Pugkhem, Alun Jones, Jordan Plieskatt, Jeremy Potriquet, Jeffery Bethony, Somchai Pinlaor, Jason Mulvenna

## Abstract

Parts of Southeast Asia have the highest incidence of intrahepatic cholangiocarcinoma (CCA) in the world due to infection by the liver fluke *Opisthorchis viverrini* (Ov). Ov-associated CCA is the culmination of chronic Ov-infection, with the persistent production of the growth factors and cytokines associated with persistent inflammation, which can endure for years in Ov-infected individuals prior to transitioning to CCA. Isobaric labelling and tandem mass spectrometry of liver tissue from a hamster model of CCA was used to compare protein expression profiles from inflammed tissue (Ov-infected but not cancerous) versus cancerous tissue (Ov-induced CCA). Immunohistochemistry and immunoblotting were used to verify dysregulated proteins in the animal model and in human tissue. We identified 154 dysregulated proteins that marked the transition from Ov-infection to Ov-induced CCA, i.e. proteins dysregulated during carcinogenesis but not Ov-infection. The verification of dysregulated proteins in resected liver tissue from humans with Ov-associated CCA showed the numerous parallels in protein dysregulation between human and animal models of Ov-induced CCA. To identify potential circulating markers for CCA, dysregulated proteins were compared to proteins isolated from exosomes secreted by a human CCA cell line (KKU055) and 27 proteins were identified as dysregulated in CCA and present in exosomes. These data form the basis of potential diagnostic biomarkers for human Ov-associated CCA. The profile of protein dysregulation observed during chronic Ov-infection and then in Ov-induced CCA provides insight into the etiology of an infection-induced inflammation-related cancer.

**Abbreviations:**

CCA: cholangiocarcinoma
Ov: Opisthorchis viverriniic
NDMA: N -nitrosodimethylamine
IHC: immunohistochemistry
MMTS: methyl methanethiosulfonate
TPP: Trans Proteomic Pipeline

## Introduction

Although a rare cancer worldwide (0.5 per 100,000 in the USA), intrahepatic cholangiocarcinoma (CCA) has the highest incidence in the world (96 per 100,000 in Northeastern Thailand [1]) in areas of the Greater Mekong subregion of Southeast Asia that overlap with transmission of the food-borne parasite *Opisthorchis viverrini* (Ov). As experimental and epidemiological evidence strongly implicate infection with this food borne pathogen in the development of CCA, Ov is one of only three eukaryotic pathogens considered Group 1 carcinogens by the International Agency for Research on Cancer (IARC) [2]. Ov infection occurs during the consumption of under-cooked fish containing the encysted metacercarial stage of the parasite [3]. After ingestion of metacercaria, the parasites excyst and migrate to the intrahepatic bile duct to mature and remain patent for decades, causing prolonged mechanical, toxicological and immunopathological damage to the biliary epithelium of the host. The pathological consequences of chronic Ov infection occur primarily in the intrahepatic bile ducts [4], where CCA also arises. The location of Ov-associated CCA makes early diagnosis difficult [5], with individuals presenting late and usually because of non-specific symptoms. Thus, despite being a slow growing tumor, Ov-associated CCA is commonly diagnosed at an advanced stage, when the primary cancer is no longer amenable to surgical extirpation and has metastasized to other organs [4]. The median survival rate for individuals after diagnosis is less than 24 months, highlighting the urgent need for diagnostic markers for Ov-associated CCA.

Although the etiology of Ov-associated CCA is multi-factorial, it is clear that the chronic inflammation and prolonged immunopathology associated with chronic Ov infection are key underlying processes in the transition to CCA [6]. As determined from our human studies in Ov endemic areas [4], Ov-associated CCA is the culmination of a series of clearly defined clinical and subclinical events caused by the persistent production of the growth factors and cytokines associated with chronic inflammation, would healing, and fibrogenesis [4,7,8]. The continuous accumulation of desmoplastic (fibrotic) elements along the intrahepatic biliary tract leads to advanced periductal fibrosis (APF) and then, in some cases, to CCA [4]. In this regard, Ov-associated CCA is not greatly different from other infection related cancers that induce a chronic inflammation that culminates in cancer: e.g., hepatocellular fibrosis and hepatocellular carcinoma from Hepatitis B and Hepatitis C viral infection [9,10]. The progression from Ov infection to CCA can be observed in a robust Syrian hamster model that uses sub-carcinogenic levels of dietary *N*-nitrosodimethylamine (NDMA) to accelerate the onset of CCA. Hamsters that are infected with Ov and fed a diet supplemented with NDMA progress to CCA while those that are infected with Ov but do not receive NDMA do not progress to CCA during a six month animal trial [11]. In this animal model of CCA, the biliary epithelium of the hamster is markedly inflamed and displays fibrosis advancing along its length after 12 weeks of infection [12], with this is fibrotic deposition routinely present at the biliary epithelium that leads to CCA [8,13]

In the current study, isobaric labelling (iTRAQ) and tandem mass spectrometry (MS/MS) were used to identify differentially expressed protein markers of CCA using animal and human models of CCA. In the hamster model of Ov-induced CCA, protein expression levels in the livers from Ov-infected hamsters that had progressed to CCA were compared to normal uninfected hamsters as well as to hamsters that were ‘Ov-infected’ but did not receive NDMA and were thus CCA-free (**Fig.** 1). Specifically, we attempted to identify protein markers of CCA that were distinct from those associated with the strong inflammation caused by Ov infection; that is, differentially expressed proteins that were either (1) associated with Ov-induced CCA but not Ov-associated inflammation when compared to normal controls, or (2) proteins associated with both inflammation and CCA but which exhibited significantly different expression in Ov-induced CCA when compared to Ov-associated inflammation. To verify these observations in human Ov-associated CCA, immunoblotting experiments and immunohistochemical analysis of CCA tissue microarrays (TMAs) were also assayed for the expression levels for these dysregulated proteins. The results of this study provide protein “signatures” of both Ov-associated inflammation and Ov-associated CCA that can be further exploited for clinical application as well as insight into the relationship between Ov-associated chronic inflammation and Ov-associated CCA.

**Figure 1:**
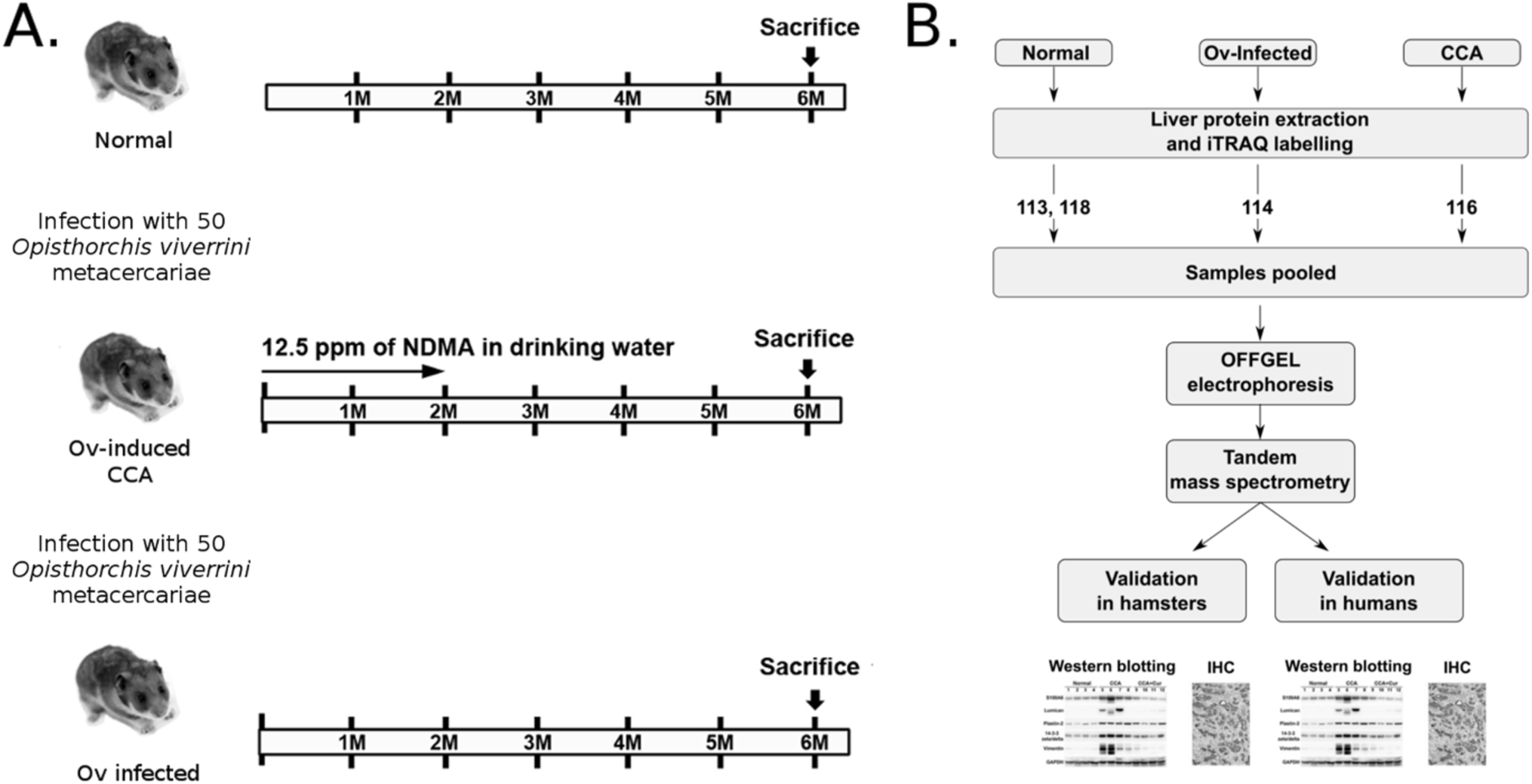
Workflow used to identify potential markers for cholangiocarcinoma. (**A**) A hamster model of CCA was used to identify dysregulated proteins in hamsters with Ov-induced CCA and in hamsters infected with Ov (‘inflammed’ state). To cause experimental Ov-induced CCA (the ‘Ov-induced CCA’ group), hamsters were infected with Ov metacercariae and had their diet supplemented with NDMA. The ‘Ov-infected’ group were infected with Ov but received no NDMA; (**B**) Hamster livers were resected and iTRAQ and tandem mass spectrometry used to profile protein expression levels. Antibodies for five proteins were then used to validate iTRAQ findings in both hamster and human tissue using immunoblotting and immunohistochemistry.

## Experimental Procedures

### Experimental design and Statistical Rationale

Quantitative iTRAQ experiments were conducted on whole liver protein preparations from three groups of hamsters: ‘Normal’, ‘Ov-induced CCA’ and ‘Ov-infected’ groups using three biological replicates from each group. For verification in hamster liver tissue western blotting experiments and immunohistochemical analysis were performed using four biological replicates from each of the three experimental groups. All human CCA tissue was obtained with informed consent from patients at Srinagarind Hospital, Khon Kaen University, Thailand. For western blotting experiments in human CCA tissue, seven biological replicates were used and in each replicate adjacent non-tumor tissue was used as a control. Candidate proteins were further verified using IHC analysis on a TMA containing tissue from 68 CCA patients, 48 male and 20 female. The aim of this study was to identify potential leads for further study and sample numbers were predominantly guided by 1) reagent costs; and 2) sample availability.

### Parasite and animal tissue

Ov metacercariae were obtained from naturally infected cyprinoid fish in Khon Kaen province, Thailand using established methods [8]. In brief, fish were digested with 0.25% pepsin-HCl and metacercariae isolated from the resulting slurry. Metacercariae were examined microscopically, counted and viable cysts were used to infect hamsters (*Mesocricetus auratus*). All animal tissue samples were taken from hamsters used in an immunohistochemical study of Ov-induced CCA described in [8] and approved by the Animal Ethics Committee of Khon Kaen University, Thailand (AEKKU 22/2557). The experimental design is shown in Fig. 1A. Hamsters were maintained at the animal research facility of the Faculty of Medicine, Khon Kaen University using protocols approved by the Khon Kaen University Animal Ethics Committee.

### Ov-induced CCA in Syrian golden hamsters

Twelve male Syrian golden hamsters (*Mesocricetus auratus*) aged between 4-6 weeks were randomly divided into three groups of four animals: 1.) the ‘Normal’ group, which received a conventional murine diet (CP-SWT, Thailand); 2.) the ‘Ov-induced CCA’ group, which were infected with 50 Ov metacercariae by oral inoculation and fed the control diet supplemented with NDMA for the first two months of the trial (administered in water available *ad libitum* at 12.5 ppm); and 3.) the ‘Ov-infected’ group, which were infected with 50 Ov metacercariae by oral inoculation and fed the control diet.

### Patient tissue

Human liver tissues were prepared as described in [14]. Written informed consent was obtained from 68 CCA patients, 48 male and 20 female, who underwent liver resection at Srinagarind Hospital, Khon Kaen University, Thailand between 1999-2010 (HE571283). Patients had a mean age in years of 57±7.7 (38-74 years). The Human Research Ethics Committee, Khon Kaen University, approved the study protocols for obtaining liver samples from the biobank of the Liver Fluke and Cholangiocarcinoma Research Center (HE571294). Frozen liver tissue from seven paired tumor cases (adjacent non-tumor and tumor) were used for western blotting and paraffin-embedded liver tissues from the same cases were used for immunohistochemistry analysis. Tissue microarrays (TMAs) were constructed by the Department of Pathology, Faculty of Medicine, Khon Kaen University as described previously [15]. The TMAs contained 68 human CCA cases. To confirm the presence of intact tumor tissue, an H&E stained section of the TMA block was prepared and reviewed by two independent pathologists. Diagnosis of CCA patients was evaluated by clinical data, imaging analysis, tumor markers, and pathology.

### Protein purification from hamster livers

Hamster liver tissue (100 mg) from each hamster in each group was suspended in 600 *µ*l of lysis buffer (7M urea, 2M thiourea, 4% (w/v) CHAPS and 40 mM Tris-Base) and homogenized with a Microtube Bead Homogenizer at 4°C for 5 min. The sample was sonicated for 1 min and incubated on ice for 30 min. The solubilized samples were centrifuged at 12,000×g for 20 min at 4°C. Protein samples were precipitated with 6 volumes of cold acetone at -20°C overnight. Proteins were then pelleted using centrifugation at 8,000×g at 4°C for 10 min and the pellets resuspended in 0.5M triethylammonium bicarbonate (TEAB) pH 8.5 (Sigma-Aldrich) and 0.1% SDS. Proteins were quantified by Bradford protein assay (Bio-Rad) using the manufacturers recommendations.

### Reduction, alkylation and iTRAQ labelling

Total liver proteins from three biological replicates were analysed in three iTRAQ experiments. Liver proteins (100 *µ*g) from three hamsters in each group were reduced, alkylated and labelled using the iTRAQ 8PLEX Multiplex Kit (AB Sciex, USA). Briefly, protein samples were reduced with 10 mM dithiothreitol at 60°C for 1 h and alkylated in 50 mM iodoacetamide or methyl methanethiosulfonate (MMTS) at 37°C for 30 min. Proteins were then digested with 2*µ*g of trypsin at 37°C for 16 h. iTRAQ labelling reagents were prepared by adding 50 *µ*l of isopropanol to each vial and these were then used to label the peptide samples for 2 hrs at room temperature. Labelled peptides were combined into three mixtures, representing three biological replicates, each containing four peptide samples (‘Ov-induced CCA’, ‘Ov-infected’ and two control channels). After labelling, peptide mixtures were cleaned using HiTrap ion exchange columns (GE Healthcare) and desalted using a Sep-Pak Vac C18 cartridge (Waters, USA). Cleaned fractions were then lyophilized prior to OFFGEL^™^.

### OGE fractionation

The 3100 OFFGEL Fractionator and OFFGEL kit pH 3–10 (Agilent Technologies, Germany) with a 24-well setup were prepared according to the manufacturer’s protocols. Lyophilized peptide mixtures were diluted to a final volume of 3.6 ml using the OFFGEL peptide sample solution. IPG gel strips (24 cm) with a 3–10 linear pH range (GE Healthcare, Germany) were rehydrated with the Peptide IPG Strip Rehydradation Solution according to the manufacturers protocol and 150 *µ*l of sample was loaded in each well. Peptides were isoelectrically focused with a maximum current of 50 *µ*A until 50 kV-h were achieved. Twenty-four fractions were recovered from each well and the wells were rinsed with 150 *µ*l of water/methanol/formic acid (49/50/1) for 15 min. Each fraction was lyophilized and then resuspended in 15 *µ*l of H_2_O with 5% (v/v) formic acid prior to LC-MS/MS analysis.

### Isolation and purification of exosomes from KKU055 cell line

The human CCA cell line KKU055 (moderately differentiated) was cultured in RPMI-1640 (GIBCO, Life Technologies) media containing 1% Penicillin-Streptomycin (GIBCO, Life Technologies) and 10% Fetal Calf Serum (GIBCO, Life Technologies) in a 25 cm^2^ culture flask (Greiner Bio One) for the first generation. For the second generation heat treated 10% FCS was used. Cell growth was carried out at 37°C under 5% CO2 and 95% humidified air. Upon reaching 80% confluency the cells were washed with 1x PBS and trypsinised using 0.25% Trypsin-EDTA (GIBCO, Life Technologies) and pelleted at 800 x g for 5 minutes before they were split into 175 cm^2^ culture flask (Greiner Bio One) in equal volumes. Approximately 800 ml of culture supernatant was collected at 80% confluency from each generation of the cell lines. Cell culture supernatants were subjected to differential high-speed centrifugation for isolation of exosomes as follows: the supernatant was centrifuged for 30 min at 2,000 x g; the supernatant was then transferred to new tubes and centrifuged for 45 min at 15,000 x g. The supernatant was then collected and centrifuged at 100,000 x g for 18 h in 3.5 25x89 mm thinwall pollyallomer tubes (Beckman) with a SW31Ti Ultracentrifuge rotor. The pellet was then resuspended in PBS, sterile-filtered (0.2 *µ*m) and centrifuged for a further 2 hours at 100,000 x g in a TLA100.3 Ultracentrifuge rotor. All centrifugation steps were performed at 4°C in order to maintain exosome stability. The final pellet was resuspended in Dulbeco’s PBS (dPBS, Life Technologies, Australia). Isopycnic separation was used to further purify exosomes using sequential ultracentrifugations in an OptiPrep (Sigma-Aldrich) iodixanol gradient as previously described [16]. Briefly, after high-speed centrifugations exosome extracts were subjected to a 0.25 M sucrose and iodixanol density gradient (5–40%). Gradients were prepared in Beckman Coulter Polyallomer 14x89mm thin-wall tubes, under sterile conditions. A volume of 200 *µ*L of exosome sample was added to each gradient column and centrifuged for 18 h, at 4°C and 100,000 x g in a SW41Ti Ultracentrifuge rotor. After centrifugation, 12x1 mL fractions were collected from each gradient column and diluted 1:3 with DPBS followed by a 2 hr centrifugation at 100,000 x g in a TLA100.3 Ultracentrifuge rotor. The pellet was then resuspended in 200 *µ*L DPBS and centrifuged for a further 30 min. Finally, the pellet was resuspended in 30 *µ*L of Tris-HCL solution (pH 7.5). Samples were stored at -80°C [17].

### Assessment of exosome presence and purity

Electron microscopy was used to identify fractions containing exosomes and to confirm the purity of exosomes in exosome preparations. A 2 *µ*l aliquot from each of the exosome samples from density gradient fractions in Tris-HCl solution (pH 7.5) was directly adsorbed onto glow-discharged formvar-carbon coated copper grids on SuperFrost slides (AgarScientific, UK) to obtain a monolayer of exosomes for analysis. Each slide was then negatively stained with freshly prepared 2 % uranyl acetate in aqueous suspension. Grids were air-dried for 3 min and imaged using a JEM-100CX transmission electron microscope (JEOL, Japan) equipped with a thermionic tungsten filament and operated at an acceleration voltage of 100 kV. Digital images were taken with a pixel size of 0.3 nm using an Olympus Morada camera using exposure times between 100 and 400 mS.

### Exosome protein purification and SDS-PAGE

Isolated exosomes were resolubilised in 1 ml 1xPBS and centrifuged at 100,000xg for 60 min at 40°C to pellet the exosomes. The pellet was resuspended in 200 *µ*l of ice cold Exosome Resuspension buffer (Total Exosome RNA and Protein Isolation Kit, Invitrogen). The sample was incubated for 5–10 min at room temperature to allow the pellet to dissolve. This was followed by gently pipetting of the sample to ensure that the pellet was completely dissolved. Acetone precipitation was carried out by adding 1:5 volume of acetone and incubating overnight at -20°C. The samples were then centrifuged at 3000 rpm for 15 min at 40°C. The supernatant was discarded and the pellet was washed again with acetone and centrifuged at 10,000 rpm for 15min at 40°C. The pellet was dissolved in 10 *µ*l of laemmli buffer and incubated at 96°C for 3–5 min. This solution was loaded on the gel against the ladder (Precision Plus Protein^TM^ Dual Colour Standards) sequence and subjected to SDS-PAGE and in-gel digestion.

Exosome protein samples were applied to 1-mm-thick 4% stacking, 12% resolving gel for SDS-PAGE. Electrophoresis was carried out at 100V for 20 minutes and then 200V for 50 minutes. The gels were stained using Coomassie Brilliant Blue and destained in 25:10:65 methanol/acetic acid/water (v/v/v). SDS-PAGE gel lanes were divided into approximately 24 slices and each slice cut into small pieces. Each gel slice was processed independently and was firstly destained twice by incubation in 50% acetonitrile, 200 mM NH_4_HCO_3_ for 45 min at 37°C and then dried using a vacuum centrifuge. The gel pieces were resuspended in 20 mM dithiothreitol (DTT) and reduced for 1 h at 65°C. DTT was removed, and the samples alkylated by the addition of 50 mM iodoacetamide and incubation in darkness at 37°C for 40 min. Gel pieces were washed twice in 25 mM NH_4_HCO_3_ for 15 min and completely dried in a vacuum centrifuge. Gel pieces were rehydrated with 20 *µ*l of trypsin reaction buffer (40 mM NH_4_HCO_3_, 10% acetonitrile) containing 20 *µ*g/ml trypsin (Sigma) for 20 min at room temperature. An additional 50 *µ*l of trypsin reaction buffer was added to the samples and incubated overnight at 37°C. The digest supernatant was removed from the gel slices, and residual peptides were washed from the gel slices by incubating three times with 0.1% formic acid for 45 min at 37°C. The original supernatant and extracts were combined and dried in a vacuum centrifuge. The tryptic peptides were resuspended in 12 *µ*l 5% formic acid before mass spectral analysis.

### Tandem mass spectrometry

Labelled peptides from iTRAQ experiments and tryptic peptides from in-gel digests of exosome proteins were analyzed by LC-MS/MS on a Shimadzu Prominance Nano HPLC (Japan) coupled to a Triple TOF 5600 mass spectrometer (ABSCIEX, Canada) equipped with a nano electrospray ion source. Two *µ*l of peptide sample was injected onto a 50 mm x 300 *µ*m C18 trap column (Agilent) at 20 *µ*l/min. The samples were de-salted on the trap column for 5 minutes using 0.1% formic acid (aq) at 20 *µ*l/min. The trap column was then placed in-line with the analytical nano-HPLC column (150 mm x 75 *µ*m C18, 5 *µ*m, Vydac, USA) for mass spectrometry analysis. A linear gradient of 1-80% solvent B (90/10 acetonitrile/0.1% formic acid (aq)) over 120 min at 800 nL/minute flow rate, followed by a steeper gradient from 40% to 80% solvent B in 5 min, was used for peptide elution. The ionspray voltage was set to 2000V, declustering potential 100V, curtain gas flow 25, nebuliser gas 1 (GS1) 10 and interface heater at 150°C. 500 ms full scan TOF-MS data was acquired followed by 20 x 50 ms full scan product ion data in an Information Dependant Acquisition (IDA) mode. Full scan TOF-MS data was acquired over the mass range 350-1800 and for product ions 100-1800. Ions observed in the TOF-MS scan exceeding a threshold of 100 counts and a charge state of +2 to +5 were set to trigger the acquisition of product ion spectra for a maximum of 20 of the most intense ions. The data was acquired and processed using Analyst TF 1.5.1 software (ABSCIEX, Canada). The MS proteomics data have been deposited to the ProteomeXchange Consortium (http://proteomecentral.proteomexchange.org) via the massIVE partner repository with the data set identifier PXD002818.

### Spectral searches and bioinformatic analysis

For iTRAQ analyses, searches were performed using ProteinPilot v4 (ABSCIEX) using the following parameters: allowing for methionine oxidation as a variable modification, carbamidomethylation (or modification with MMTS where appropriate) as a fixed modification, two missed cleavages, charge states +2, +3 and +4 and trypsin as the enzyme. Searches were conducted against the Uniprot hamster proteome dataset (*Cricetulus griseus*; proteome id UP000001075) as of Oct 2013 (23,878 entries). Proteins were grouped using ProteinPilot’s ProGroup algorithm, single peptide identifications were not considered and only proteins containing at least one unique, significant peptide identification were reported. Searches were also conducted with X! TANDEM Jackhammer TPP (2013.06.15.1) [18] using the same database appended with reversed sequences with the following parameters: enzyme = trypsin; precursor ion mass tolerance =± 0.1 Da; fragment ion tolerance =± 0.1 Da; fixed modifications = carbamidomethylation (or modification with MMTS where appropriate) and iTRAQ modification of Lys and N-term free-amines (using modification masses 304.205360 and 304.199040); variable modifications = methionine oxidation and variable labelling of Tyr residues with iTRAQ reagents; number of missed cleavages allowed = 2; allowed charge states = +2–+4; and ‘k-score’ as the scoring algorithm. The Trans Proteomic Pipeline (TPP) [18] was used to validate peptide and protein identifications using PeptideProphet [19] and ProteinProphet [20] and Mayu [21] was used for false discovery rate (FDR) estimation. Using the TPP and the same parameters, with the exception of Lys and N-terminal modification by iTRAQ reagent specified as a variable modification, further searches were conducted to estimate the iTRAQ labelling efficiency. GO enrichment was performed using the BiNGO plugin [22] in Cytoscape [23]. Hamster proteins were mapped to their corresponding mouse protein using BLASTp and BiNGO used to identify enriched GO terms using the *Mus musculus* annotations and default parameters. iTRAQ reporter ion intensities from peptides identified by ProteinPilot as suitable for quantitation and which possessed a probability greater than 0.95 (Suppl. Table 6) were used in the R package iQuantitator [24] to generate credible intervals for protein expression differences in specific samples. Filtering of peptides for quantitation included the removal of any peptide without the “auto” annotation, including those peptides with “auto - shared MS/MS” providing a set of peptides uniquely assigned to protein identifications. iQuantitator uses a model-based approach to describe variation in observed data and Bayesian inference to estimate credible intervals for protein expression across multiple iTRAQ experiments and proteins were considered up, or down regulated if the start and end of computed 95% credible intervals were >1, or <1 respectively. Only proteins quantified on the basis of two significant peptides were considered. All data and search results were submitted to ProteomeXchange with the accession number PXD002818. Spectral data from the analysis of exosome proteins were searched against the UniProt human proteome database (as of the 1st of February, 2016; 70,611 sequences) using the TPP as described above but without iTRAQ labelling specified as modifications. To determine which exosome proteins were homologous to hamster proteins identified as significantly dysregulated, the ‘blastp’ program was used to search human proteins against a database of hamster proteins with a bit score cutoff of 50.

### Western blotting

For immunoblotting of hamster and human liver tissue, 120 mg of liver tissue was minced and incubated on ice for 30 min in ice-cold lysis buffer after which the protein concentration was determined using the Bradford assay. Ten micrograms of liver protein was separated on a 12% SDS-PAGE gel and transferred to a polyvinylidene difluoride membrane (PVDF, Amersham Bioscience, Piscataway, NJ, USA) for 2 h at 60 V. The membrane was incubated overnight at 4°C with primary antibody, diluted in 2% nonfat dried milk/ 1X Phosphate Buffered Saline Tween-20 (PBST). The membrane was then incubated with the appropriate horseradish peroxidase-conjugated secondary antibody (1:3,000, GE healthcare) diluted in 2% nonfat dried milk/PBST and visualized by enhanced chemiluminescence using ECL Western blotting Detection Reagent (GE Healthcare). The ImageQuant TL software v2005 (1.1.0.1) (Non-linear Dynamics, Durham, NC) was used for quantitative analysis of each band. The relative band intensity of the treated group was normalized by average normal control to adjust for experimental variation.

### Immunohistochemistry and scoring

Hamster tissue (n=4 from each group), human paired tumor and distal normal tissue (n=7) and human TMAs (n=68) were cut into 5-*µ*m sections and mounted on silane-coated slides (Sigma, St. Louis, MO, USA) using the immunoperoxidase method [25]. Sections were deparaffinized in xylene and hydrated in graded alcohols. For antigen retrieval, slides were floated on 10 mM citric acid buffer (pH 7.0) and heated for 10 min at 110°C and allowed to cool. Slides were washed and blocked for endogenous peroxidase in 3% hydrogen peroxide diluted in PBS for 15 min. After washing with PBS, sections were blocked with 5% fetal bovine serum for 30 min and then incubated at 4°C overnight with primary antibodies. Sections were washed with PBS and incubated with the appropriate secondary antibodies for 30 min at room temperature. Sections were then developed with 3,3’-diaminobenzidine (DAB; Sigma), counter-stained with Mayer hematoxylin and mounted in Permount. The staining density and intensity in hamster tissue was scored as described previously [26]. Briefly, the intensity of protein expression was graded as follows: 0, no staining; 1+, mild; 2+, moderate; and 3+, strong. The staining density was quantified as the percentage of cells stained positively in tissue as follows: 0, no staining; 1, positive staining in <25%; 2, 25–50%; and 3, >50%. The intensity score was multiplied by the density score to yield an overall score of 0–9 for each sample. For hamster and human tissue, all sections were evaluated by three investigators, blinded to grading in human and hamster sections, and when there was consensus between two of the three investigators this grade was designated as acceptable. For human tumor tissue in TMA experiments, 68 cases provided triplicate tissue spots and positive areas were categorized as follows: negative (<10%) and positive (≥10%) as described previously [27]. As described above, all sections were evaluated by three investigators blinded to grading and the consensus of two from three investigators was designated an acceptable grade.

### Statistical analyses

In western blot analysis protein expression levels in each group were compared and statistical significance of band intensity was assessed using the Student’s t-test and the data expressed as mean ± S.D. For statistical analysis of IHC experiments in hamster tissue a nonparametric Mann-Whitney U test was used to compare the IHC expression level of each protein. Statistical analyses were performed using SPSS version 15 (SPSS, Inc, Chicago, IL). A *p*-value of less than 0.05 was considered statistically significant.

## Results

### O. viverrini induced CCA in hamsters

The experimental design is set out in Fig. 1A; briefly, three groups each consisting of four hamsters (N = 4) were maintained for six months as follows: an ‘Ov-induced CCA’ group with hamsters proceeding to Ov-induced CCA (i.e. diet supplemented with NDMA); an ‘Ov-infected’ group, with hamsters infected with Ov but without CCA (i.e. diet not supplemented with NDMA); and a ‘Normal’ group with hamsters without Ov-infection or Ov-induced CCA. For iTRAQ experiments, total protein from livers from three hamsters in each group were used as biological replicates (Fig. 1B).

### Quantification of protein expression in hamster livers

Trypsinized and isobarically labelled total hamster liver protein, in triplicate, were fractionated using OFFGEL electrophoresis (OGE) and subjected to tandem mass spectrometry. ProteinPilot searches of the UniProt hamster proteome database identified 904 proteins with at least two significant peptides and an Unused score >= 2.3 (corresponding to a *p* -value cutoff of <0.01) (Suppl. Table 1a). Local false discovery rates, calculated using the nonlinear fitting method [28], were below 1% for each replicate. Similar results were obtained (769 identified proteins; at least two unique peptides and a FDR of < 0.01) using the TPP and Mayu for FDR estimation (Suppl. Table 1b). The iTRAQ reporter ions from each label were used to quantitate protein levels in the experimental groups (for an example spectra see Fig. 2A). Labelling efficiency was assessed in searches conducted using iTRAQ labelling as a variable modification and 96% of Lys residues in peptides confidently (*p* < 0.05) assigned to proteins were modified with an iTRAQ label. Approximately 80% of the N-termini were labelled, the lower percentage most likely due to N-terminal modification or blocking and/or peptide degradation after labelling [29]. Peptides with iTRAQ labelled Tyr residues accounted for only 0.02% of identified peptides and these were excluded from quantitative calculations (Fig. 2B). The comparison of the two control channels showed that 95% of the identified proteins had expression ratios within 1.2 and 0.86 while control versus test samples exhibited much greater variation (Fig. 2C).

**Figure 2:**
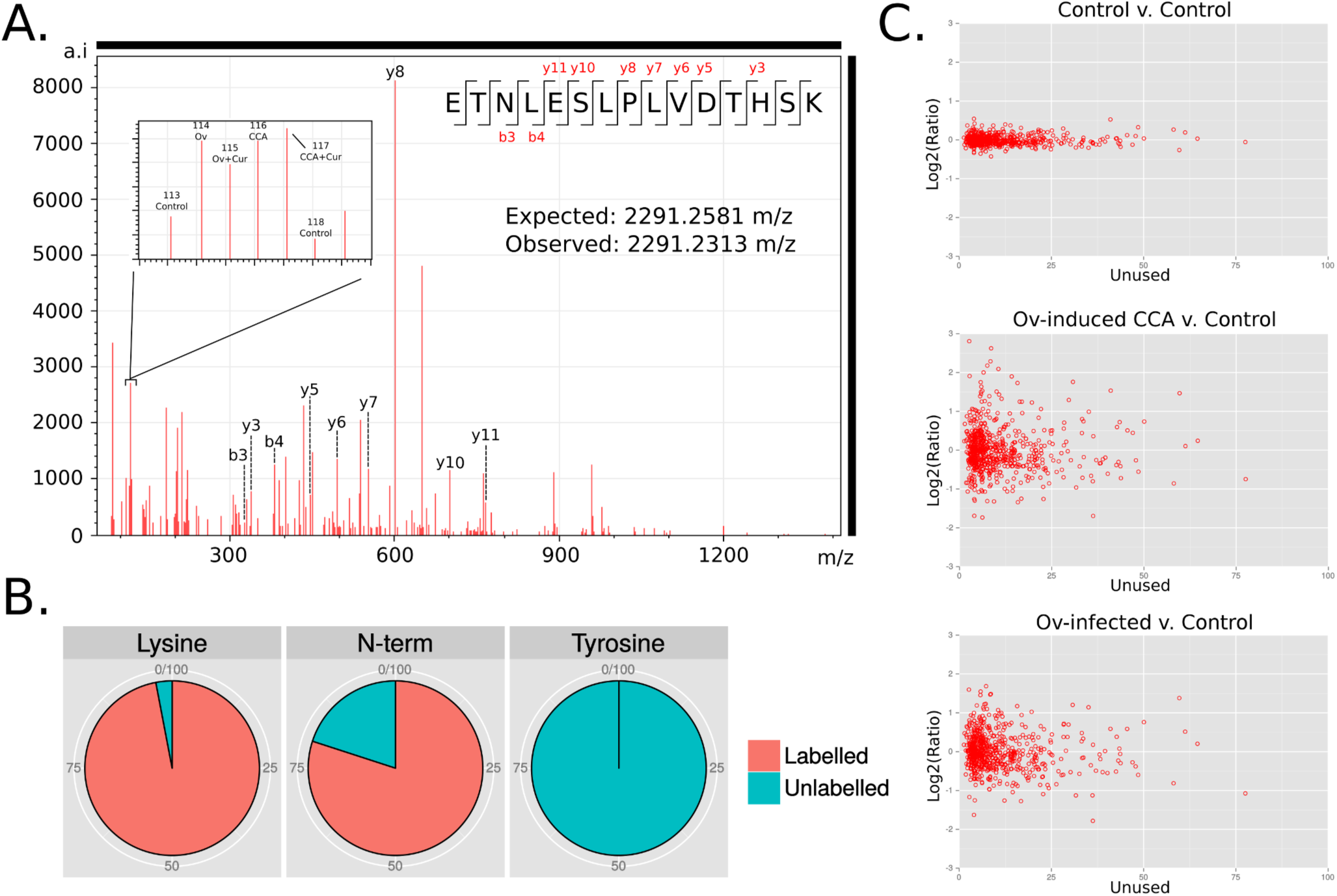
Isobaric labelling (iTRAQ) of hamster liver proteins. (**A**) Representative spectrum from tandem mass spectrometry of hamster liver proteins with reporter ions used for quantitation (inset); (**B**) Labelling efficiency was assessed using searches specifying iTRAQ labels as variable modifications; (**C**) In each iTRAQ replicate two aliquots of the same control sample were labelled with different reporter ions and used as internal validation of expression ratios.

### Proteins dysregulated during Ov infection and Ov-induced CCA

The program iQuantitator [24] was used to identify significantly dysregulated proteins in the ‘Ov-induced CCA’ and ‘Ov-infected’ groups compared to the ‘Normal’ group. In the ‘Ov-induced CCA’ group 246 proteins (137 under-expressed and 109 over-expressed) were dysregulated and, in the ‘Ov-infected’ group, 121 proteins (69 under-expressed proteins and 52 over-expressed) were dysregulated. A comparison of dysregulated proteins in these two groups further showed that 92 proteins were significantly dysregulated in both the ‘Ov-induced CCA’ and the ‘Ov-infected’ groups when compared to the ‘Normal’ group, while 154 proteins were dysregulated only in the ‘Ov-induced CCA’ group and 29 were significantly dysregulated only in the ‘Ov-infected’ group (Fig. 3, Suppl. Table 2 and 3). GO enrichment analysis of proteins dysregulated in the ‘Ov-induced CCA’ group showed enrichment (*p* < 0.05) for metabolic and catabolic “biological process” terms typically associated with cancer, including organonitrogen compound biosynthetic processes and the metabolism and catabolism of hydrogen peroxide, as well as terms related to the response to ‘wound healing’ and ‘cell-cell adhesion’ (Suppl. Table 4a). GO enrichment analysis of protein dysregulated in the ‘Ov-infected’ group showed a very similar profile of enriched ‘biological processes’ with all 40 terms enriched in this group also enriched in the ‘Ov-induced CCA’ group (Suppl. Table 4b). Of the 92 proteins identified as dysregulated in both the ‘Ov-infected’ and ‘Ov-induced CCA’ groups, when compared to the ‘Normal’ group, all exhibited a pattern of expression in which the ‘Ov-induced CCA’ group showed the greatest magnitude of dysregulation when compared to the ‘Ov-infected’ group. When the ‘Ov-induced CCA’ and ‘Ov-infected’ groups were directly compared, 165 proteins were identified as dysregulated (77 down-regulated and 89 up-regulated; Suppl. Table 2c), including 50 of the 92 proteins that had been identified as dysregulated in both experimental groups when compared to the normal controls. Many of these 50 proteins, for example vimentin and annexin A1 [30], have been associated with different malignancies which, when combined with the very similar enriched GO terms from the ‘Ov-induced CCA’ and ‘Ov-infected’ groups, suggests that some protein expression changes associated with cancer are occurring during infection and that increased expression of these proteins could mark the transition from infection to CCA.

**Figure 3:**
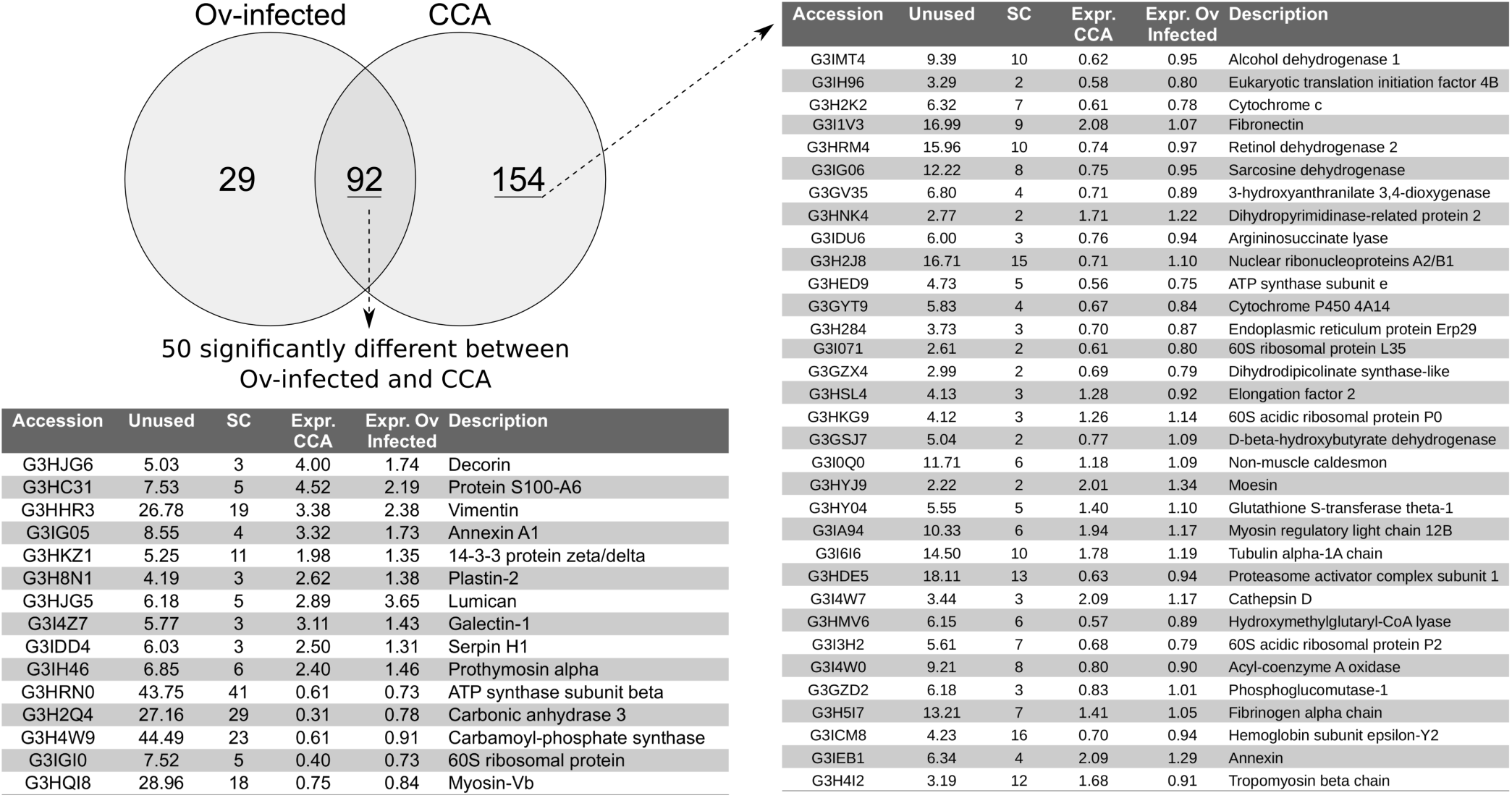
Dysregulated proteins common to CCA and Ov-infected group. Selected proteins identified as dysregulated in the livers of hamsters with experimentally-induced CCA and hamsters infected with Ov. Abbreviations used: Unused — ProteinPilot score for protein identification; SC — number of unique peptides used for protein identification; Expr. CCA — fold-change of protein when compared to normal control sample; Expr. Ov infected — relative fold-change when compared to normal control sample.

### Validation of iTRAQ data in hamster tissue

To provide an independent measure of protein expression in hamster tissue five proteins were selected for immunohistochemistry (IHC) and immunoblotting experiments on the basis of 1.) the strength of their dysregulation and 2.) the functional significance of the protein. In the ‘Ov-induced CCA’ group, immunoblotting of five proteins (S100A6, lumican, plastin-2, 14-3-3 zeta/delta and vimentin) exhibited the same expression profile observed in iTRAQ experiments (Fig. 4A and B). In the ‘Ov-infected’ group, four proteins (prolargin, plastin-2, 14-3-3 zeta/delta, and vimentin) also exhibited the same expression profile in immunoblotting experiments as was observed in iTRAQ experiments (data not shown). In IHC experiments using graded tumor tissue from the ‘Ov-induced CCA’ group, expression of S100A6, lumican, plastin-2 and 14-3-3 zeta/delta was observed in the cytoplasm of bile duct tumor, while vimentin was found mainly in the cytoplasm of fibroblasts at periductal fibrosis and tumor stroma (Fig. 4C). Once again, protein expression mirrored that observed in the iTRAQ experiments (Fig. 4C and D).

**Figure 4:**
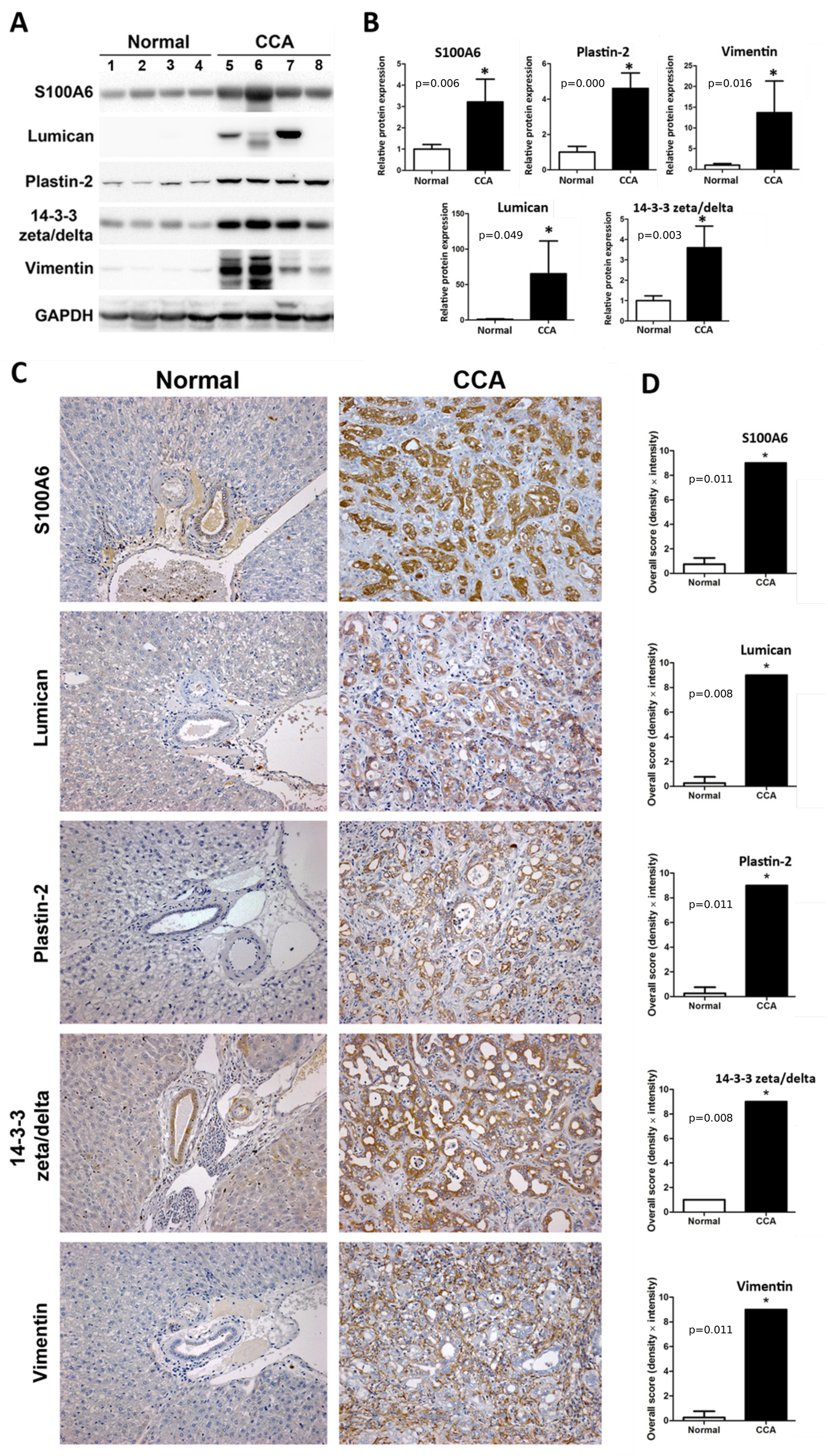
Validation of iTRAQ data in hamster tissue. Expression changes of five proteins (S100A6, lumican, plastin-2, 14-3-3 zeta/delta and vimentin) were evaluated by western blot and Immunohistochemistry (IHC). (**A**) Western blot analysis of candidate proteins in normal (lane 1-4) and ‘Ov-induced CCA’ groups (lane 5-8); (**B**) The relative band intensity of the western blot in the ‘Ov-induced CCA’ group was normalized by the average normal and the results are shown as a bar graph; (**C**) A representative immunohistochemistry experiment showing staining of five proteins in hamster livers. The sample shown is representative of each group (4 cases/group; original magnification,× 200); (**D**) Protein expression levels were estimated from (**C** by intensity score and multiplied with the density score to yield an overall grading score. The grading results are shown as a bar graph. In (**B**) and (**D**), asterisk (*) denotes significant difference (*p* < 0.05) by Student’s T-test versus normal and significant *p* values are shown on bar plots.

### Validation of iTRAQ data using immunoblotting and immunohistochemistry in human tissue tissue

To determine the relevance of the hamster data to human Ov-associated CCA, five proteins (S100A6, lumican, plastin-2, 14-3-3 zeta/delta, vimentin) that were over-expressed in the ‘Ov-induced CCA’ hamster group were assayed in human CCA cases. Paired tumor and distal normal tissue (i.e tissue taken at a site near the CCA tumor but with no evidence of frank carcinoma or observed dysplasia) from seven human CCA cases were analysed using immunoblotting (Fig. 5A) with the relative band intensity of all proteins shown in Fig. 5B. Protein S100A6, 14-3-3 zeta/delta and vimentin were significantly increased in human CCA tumor tissue while lumican and plastin-2 were not significantly up-regulated, although there was a trend toward increased expression in the latter two proteins. IHC analysis of the same seven samples showed that, as in the hamster experiments, S100A6, lumican, plastin-2, 14-3-3 zeta/delta were observed in the cytoplasm of bile duct tumor, while vimentin was found mainly in the cytoplasm of fibroblasts at periductal fibrosis and tumor stroma (Fig. 5C). Tissue adjacent to tumor tissue but with no evidence of frank carcinoma showed only minor staining (Fig. 5C) in all seven cases. To examine protein expression in a larger number of samples we used IHC analysis of a human CCA TMA to investigate positive staining of the five proteins used in the immunoblotting experiments in 68 human CCA cases. Positive staining rates in tumor tissues were 78% (53 of 68) for S100A6, 84% (57 of 68) for lumican, 50% (34 of 68) for plastin-2, 82.4% (56 of 68) for 14-3-3 zeta/delta and 98.5% (67 of 68) for vimentin.

**Figure 5:**
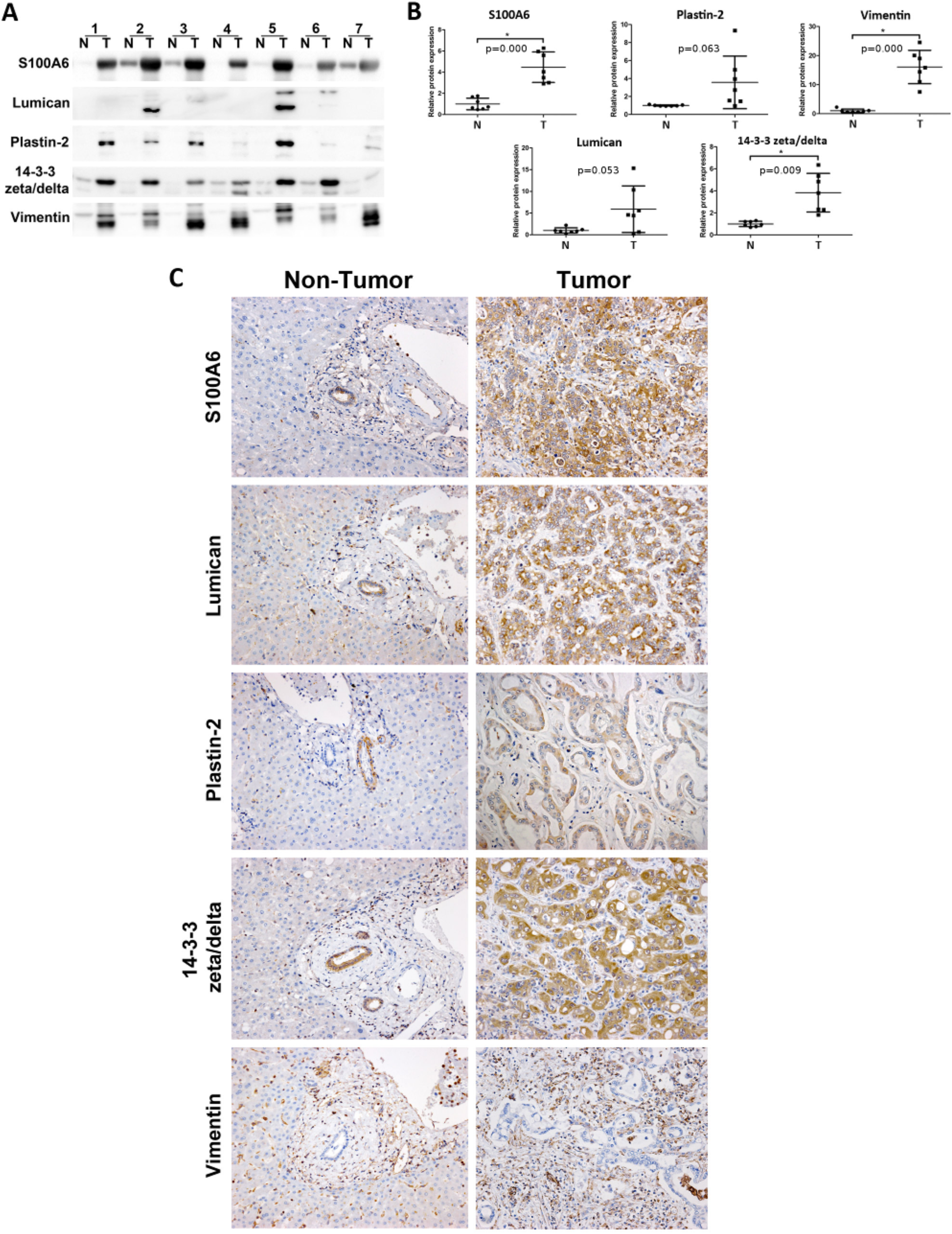
Validation of iTRAQ data in human tissue. Expression changes of five proteins (S100A6, lumican, plastin-2, 14-3-3 zeta/delta and vimentin) were evaluated by western blot and immunohisto-chemical staining of human tissue. (**A**) Western blot analysis in seven paired tumor cases (N, adjacent non-tumor and T, tumor tissue); (**B**) the relative band intensities were normalized by average normal and plotted as a scatter plots. Asterisk (*) denotes significant difference (*p* < 0.05) by Student’s T-test versus ‘Normal’ group and *p* -values are shown on the bar plots; (**C**) Immunohistochemical staining of the five proteins in the same human tumor tissue samples showed only minor staining in non-tumor tissue (original magnification,× 200).

### Identification of potential circulating markers for CCA

A number of studies have now established that exosomes secreted by cancer cells can be differentiated on the basis of their protein cargo and that these proteins show cell-type specificity [31]. Accordingly, great interest has been shown in exploiting circulating exosomes as biomarkers of disease or as a mechanism for reducing the dynamic range of the circulating proteome [32]. In order to identify potential circulating markers of CCA, we characterized the proteome of exosomes secreted by the KKU055 human CCA cancer cell line and compared them to CCA-associated proteins identified in the hamster experiments. Tandem mass spectrometry of exosomes purified from KKU055 culture medium using differential centrifugation and density gradient centrifugation provided 282 identifications at a FDR of < 0.01 (each identification comprising of at least two peptides at least one of which was unique; Suppl. Table 7). These proteins were compared to proteins significantly dysregulated in the ‘Ov-induced CCA’ hamster group using BLAST and 27 proteins were found in common (summarized in Fig. 6 and in Suppl. Table 8). These proteins included some of the most highly up-regulated proteins identified in the ‘Ov-induced CCA’ group, including vimentin, galectin-1, vitronectin and Protein S100; two of these, vimentin and protein 14-3-3, were also validated as up-regulated in human immunoblotting and IHC experiments. The presence of these CCA-associated proteins in exosomes secreted by a human CCA cell line suggests that these proteins are important targets for future validation in CCA patients as potential diagnostic and/or prognostic protein markers.

**Figure 6:**
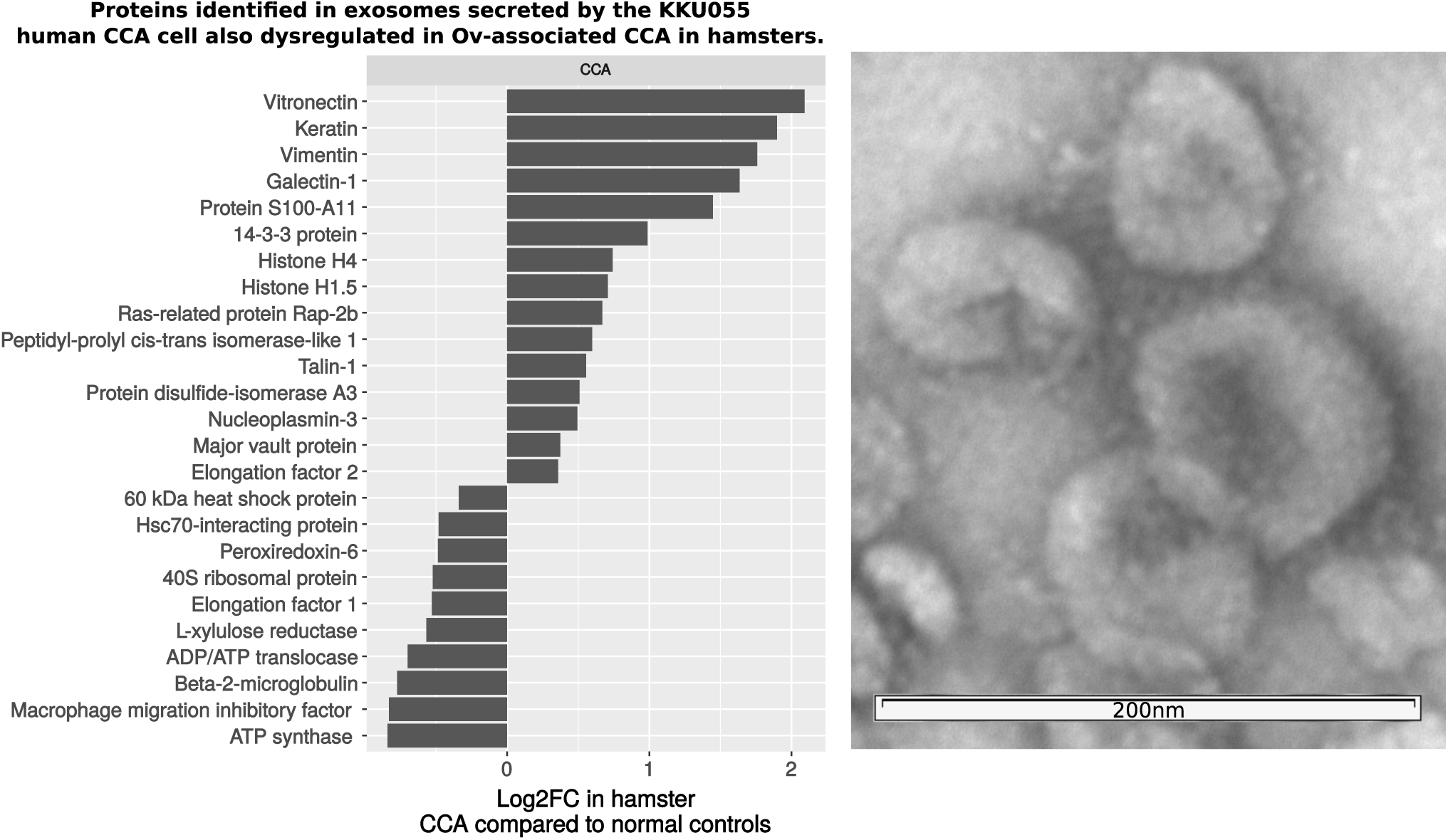
Proteins homologous to hamster proteins dysregulated in CCA that were identified in exosomes secreted by the human CCA cell line KKU055. Twenty-seven human exosomes proteins, from 282 total identifications, were found to be homologous (BLAST bit score >50) to proteins dysregulated in the hamster model of CCA. These included some of the most highly over-expressed proteins in hamster livers of hamsters with CCA. On the right, transmission electron micrograph of exosome preparation used in proteomics experiments.

## Discussion

Ov-associated CCA presents an exceptional model for investigating the role of inflammation-associated cancers, as it progresses through a series of well-defined clinical stages that can be detected in both a human and animal model. Moreover, the chronic inflammation during Ov infection, which can last for years, has been shown to have an etiologic role in the development of Ov-associated CCA in the human and animal models of this bile duct cancer. The pattern of protein expression in the ‘Ov-infected and ‘Ov-induced CCA’ hamsters groups showed protein dysregulation profiles that could discriminate between the subclinical and clinical stages leading to Ov-associated CCA. Protein markers that could discriminate between Ov infection and Ov-associated CCA are of particular interest in areas where Ov is endemic and the prevalence of Ov infection rates can reach as high as 75% of the population [4]. In our previous work, we have shown that systemic and Ov-specific interleukin-6, a known inflammatory cytokine, are associated with Ov-associated CCA and that this cytokine possesses a positive association with advanced periductal fibrosis, a precursor stage to CCA and to Ov-associated CCA itself [33]. However, the low diagnostic specificity of IL-6, which has been proposed as a inflammatory marker for many conditions (e.g. cancer [34], mechanical injury [35] and bacterial infection [36] among others) decreases its utility as a diagnostic marker for CCA. Proteins in clinical use as markers of Ov-associated CCA suffer from a similar lack of diagnostic specificity: e.g., the carbohydrate antigen (CA-19-9) is elevated in Ov-associated CCA but also in primary biliary cirrhosis and in individuals who smoke [37]. Similarly, the carcinoembryonic antigen (CEA) is elevated in only ~30% of Ov-associated CCA cases. In the current study, we used advanced proteomic techniques (isobaric labelling and tandem mass spectrometry) on animals at the distinct stages in the transition from Ov-infection to Ov-induced CCA to identify a large number of potential markers for Ov-associated CCA that could also distinguish Ov-associated CCA from the inflammation which accompanies chronic Ov-infection in humans resident in Ov endemic areas and which could be further evaluated for their application as diagnostic markers of CCA.

To identify potential markers for CCA, we identified proteins that were dysregulated in animals with CCA (‘Ov-induced CCA’ group) but not during Ov infection (‘Ov-infected’ group). Over 150 of these proteins were identified, including many proteins previously associated with cancer. In particular, proteins associated with the extracellular matrix and cell mobility and adhesion were over-expressed in the CCA group, including vitronectin [38], fibronectin [39], septin-7 [40] and talin-1 [41] all of which play key roles in the remodelling of the extra-cellular matrix (ECM), adhesion and metastasis known to occur during human CCA. Likewise, over-expression of Ras-related protein Rap-1b and serum amyloid A protein have been observed as key events during carcinogenesis in several malignancies, the former mediating malignant transformation, proliferation and cell growth [42] and the latter suppressing cell immunity via stimulation of immunospressive neutrophils to produce interleukin-10 [43]. These proteins were not dysregulated in the ‘Ov-infected’ group suggesting that proteins associated with these processes can act as markers for the onset of cancer and can discriminate infection from cancer. Likewise we identified a number of cancer-associated proteins, for example vimentin, lumican and galectin-1, that were already over-expressed during infection, suggesting that some of the key processes that become important during carcinogenesis are active during infection. If detectable in blood, or other matrices, and both vimentin [44] and galectin-1 [45] were detectable in exosomes from the KKU055 cell line, these proteins could provide useful tools for monitoring infection-related inflammation and, potentially, the transition to CCA.

Ninety-two proteins were dysregulated in both the ‘Ov-infected’ and ‘Ov-induced CCA’ groups and the majority of these were shown to be differentially expressed when the ‘Ov-infected’ and ‘Ov-induced CCA’ groups were directly compared (Fig. 3) suggesting that processes activated during the inflammatory response continue after transformation to a tumor. In particular, proteins closely associated with the pro-inflammatory transcription factor NF-κB showed dysregulation during infection (‘Ov-infected’ group) and a significantly greater magnitude of dysregulation during cancer (‘Ov-induced CCA’ group). Activation of NF-κB promotes infiltration of inflammatory cells and alterations to the redox environment. Prolonged activation can induce oxidative DNA damage via constitutive increases in reactive oxygen (ROS) and nitrogen species (NOS). NF-κB activators, e.g. clusterin and S100A10, and targets, e.g. S100A6, showed increased dysregulation in the ‘Ov-induced CCA’ group when compared to the ‘Ov-infected’ group (Fig. 3; Suppl. Table 2) as did the downstream targets of S100A6: annexin 2, tropomyosin β, annexin 11 and lamin B1. Similarly, CCA cell lines (KKU100, KKU-M156 and KKU-M213) exhibit constitutive activation of NF-κB [46] and the results presented here also suggest constitutive activation of NF-κB as an underlying cause of carcinogenesis. In the context of human Ov infection, decades of sustained injury to the bile duct epithelium by the parasite likely leads to constitutive expression of NF-κB, creating a “smouldering inflammatory milieu” [4,47] that, ultimately, creates severe hepatobiliary abnormalities, principal among them Ov-associated CCA.

On the basis of immunoblotting and IHC experiments, the protein expression profile generated from the hamster model appeared to provide a reasonable approximation of protein expression in the human CCA disease. Three of the five proteins used to verify iTRAQ results in human tissue were found to be significantly over-expressed in immunoblotting experiments and, in IHC experiments on TMA using 68 CCA cases, all but one protein (plastin-2) exhibited increased staining in at least 78% or more of tumors. Thus, not only could the three proteins form the basis of a potential marker for CCA, but these results also suggest that many of the 243 other proteins identified as over-expressed by iTRAQ may also be potential markers for human CCA. In non-endemic regions, no specific and sensitive marker for CCA is in clinical use, as discussed above, CA-19-9 and CEA suffer from a lack of clinical specificity. Several proteins have been associated with CCA in a non-Ov-associated model; both circulating apolipoprotein 1 [48] and vimentin [49] have been associated with CCA and both of these proteins were identified as over-expressed by iTRAQ in this study (Suppl. Table 3). One quantitative proteomic study of human non Ov-associated intrahepatic CCA tumor tissue identified 39 dysregulated proteins that are consistent with the dysregulated proteins reported here. Of 16 up-regulated proteins observed in that study, ten proteins were also found to be significantly over-expressed in the current study including, vimentin, protein S100-A11, profilin-1, transgelin-2, prothymosin and cofilin [49]. Likewise, the overexpression of 14-3-3 zeta/delta has recently been demonstrated in both advanced fibrotic lesion and CCA tissue showing its potential as a marker of risk for Ov-associated CCA [50]. Taken together, these similarities suggest that the potential biomarkers identified here will also have utility for the diagnosis of non-Ov-associated CCA.

To be useful, particularly in a resource-poor environment, protein markers need to be detectable in an easily accessible matrix such as blood, urine or saliva. In this work we used exosomes from the human CCA cell line KKU055 to determine which CCA-associated proteins are likely to be identifiable in the circulation of CCA patients. Proteins identified in the exosomes included human homologues of the most up-regulated proteins in the hamster model of CCA and key proteins associated with carcinogenesis such as vitronectin, talin-1 and Ras-related protein Rap-1b. These proteins are now being assessed in detail as potential circulating markers of CCA in human patients. As discrete packages of protein and RNA emanating directly from tumor cells, exosomes have a number of advantages when considered as biomarkers: exosomes protect their protein cargo from proteolytic digest; they can be purified from plasma or sera, reducing the dynamic range of this matrix; and previous studies have demonstrated that the number of exosomes in circulation increases in cancer patients [51]. Accordingly, exosomes purified from blood and urine are now being assessed as markers for a wide range of malignancies, including prostrate [52], lung [53], breast [54] and ovarian cancers [55], among others [32]. The work performed in this study now enables a targeted examination of potential protein markers of CCA in exosomes not only from blood but other matrices such as urine and saliva.

In general, cholangiocarcinogenesis is associated with a wide range of cellular changes that are reflected in the protein expression profile of tumor tissue. In the current study, we utilized a robust hamster model of Ov-induced CCA to profile these protein expression profiles using quantitative protein methods iTRAQ and tandem mass spectrometry to identify potential markers in the transition from Ov-infection to Ov-associated CCA. Using isobaric labelling, we identified over 200 dysregulated proteins that can now inform future efforts to develop a biomarker for the diagnosis of Ov-associated CCA. The over-expression of five proteins was confirmed in hamster tissue and, more importantly, three of these five proteins were then confirmed as over-expressed in human Ov-associated tumor tissue. The latter observation suggests that over-expressed proteins identified in the hamster model of Ov-induced CCA may also be over-expressed in human Ov-associated CCA. Taken together, these data provides a basis for future efforts to develop biomarkers and potential therapeutics for the diagnosis and treatment of Ov-associated CCA as well as other infection-related cancers.

## Acknowledgments

This work was supported by the Higher Education Research Promotion and National Research University Project of Thailand, Office of the Higher Education Commission through the Health Cluster (SHeP-GMS; PD55208 and NRU572004 to JK and SP), Thailand Research Fund through the Royal Golden Jubilee Ph.D. Program, Thailand (JK and SP), award R01CA155297 (JMB and JPM) from the National Cancer Institute, award P50AI098639 (JMB) from the National Institute of Allergy and Infectious Disease and fellowship support (JPM) and research support (JMB and JPM - grant number 1051627) from the National Health and Medical Research Council of Australia.

